# Targeting MondoA Sensitizes B-ALL Cells to Asparaginase and Reveals Glutathione-Associated Protection

**DOI:** 10.64898/2026.11.20.768131

**Authors:** Alissia Fernandes Madeira, Constantin Segner, Christian Brückner, Busheng Xue, Alisa Kolesnikova, Julia Mergner, Alexandra Sipol, Elmar Wolf, Roland Rad, Julia Hauer, Jürgen Ruland, Poul H Sorensen, Stefan Burdach

## Abstract

**Background:** L-Asparaginase (ASNase) is central to acute lymphoblastic leukemia (ALL) therapy, but resistance and long-term cytotoxic effects remain challenging. We investigated whether MondoA (MLXIP) modulates ASNase sensitivity.

**Methods:** Viability and apoptosis were assessed in CRISPR-Cas9-generated MondoA- knockout (MKO), MondoA-overexpressing (MOE), and Cas9-only control (WT) Nalm6 B-ALL cells following ASNase exposure or glutamine depletion. Omics, ChIP-seq, and patient-RNA- seq examined MondoA-ASNS associations.

**Results:** MondoA deficiency was associated with reduced ASNS mRNA and protein abundance. MYC binding to the ASNS promoter was reduced in MKO cells, suggesting an indirect transcriptional link. MondoA and ASNS expression correlated in patient samples, particularly in T-ALL (r = 0.769). MKO cells showed increased ASNase, but not doxorubicin or vincristine, sensitivity. Glutamine withdrawal during ASNase treatment further reduced viability. GSH-associated protection was strongest in biallelic MKO2.

**Conclusion:** MondoA status was associated with ASNS abundance and ASNase response, warranting further investigation of these processes as candidate determinants of ASNase sensitivity.

## Introduction

Acute lymphoblastic leukemia (ALL) is the most common pediatric malignancy.(1) Although five-year survival reaches approximately 80–90%, cytotoxic chemotherapy causes severe long-term late effects, including secondary malignancies, coronary disease, osteoporosis, endocrinopathies and post-chemotherapy cognitive impairment.(2) Defining mechanisms of drug resistance and improving the therapeutic index therefore remain crucial.

Asparaginase (ASNase) catalyzes the conversion of asparagine (Asn) to aspartate and, to a lesser extent, glutamine to glutamate, rendering Asn functionally essential for many B-ALL cells due to their reduced expression of asparagine synthetase (ASNS).(3) ASNase also increases mitochondrial reactive oxygen species (mtROS) and promotes apoptosis.(4, 5) Refractory leukemias can reactivate ASNS through promoter demethylation or stress-response pathways, including the Amino Acid Response (AAR) Pathway.(6)

We previously identified MondoA (encoded by MLXIP) as a malignancy factor in pediatric B-precursor ALL (B-ALL) that adapts leukemic cells to metabolic stress, limits ROS, and confers resistance to glutamine (Gln) starvation. MondoA and MYC are members of the extended MYC network, which coordinates nutrient utilization with cell growth.(7, 8) Because ASNS uses Gln as the amide donor for de novo Asn synthesis, MondoA-mediated adaptation to Gln starvation may also influence the response to ASNase-induced amino acid depletion.

We therefore hypothesized that MondoA might protect B-ALL cells against ASNase by supporting ASNS expression and adaptation to the metabolic stress caused by Asn and Gln depletion. We examined ASNS abundance and MYC promoter occupancy, ASNase sensitivity, and the effects of Gln withdrawal and reduced glutathione (GSH) supplementation.

## Materials and methods

### Cell models

Nalm6 MondoA-knockout clones (MKO1 and MKO2), a MondoA-overexpressing clone (MOE), and a Cas9-only control clone (Nalm6 + Cas9, hereafter WT) were established as previously described.(7) MKO1 has monoallelic and MKO2 has biallelic MondoA disruption. MondoA protein expression in these clones was verified by Western blotting (Figure 1A). Both knockout clones were included in ASNase viability, Gln-withdrawal, and GSH endpoint experiments. The ASNase apoptosis titration and comparator-drug experiments used MKO1; the GSH time courses compared WT with MKO2. Clone identity is specified for each experiment.

**Figure 1:**
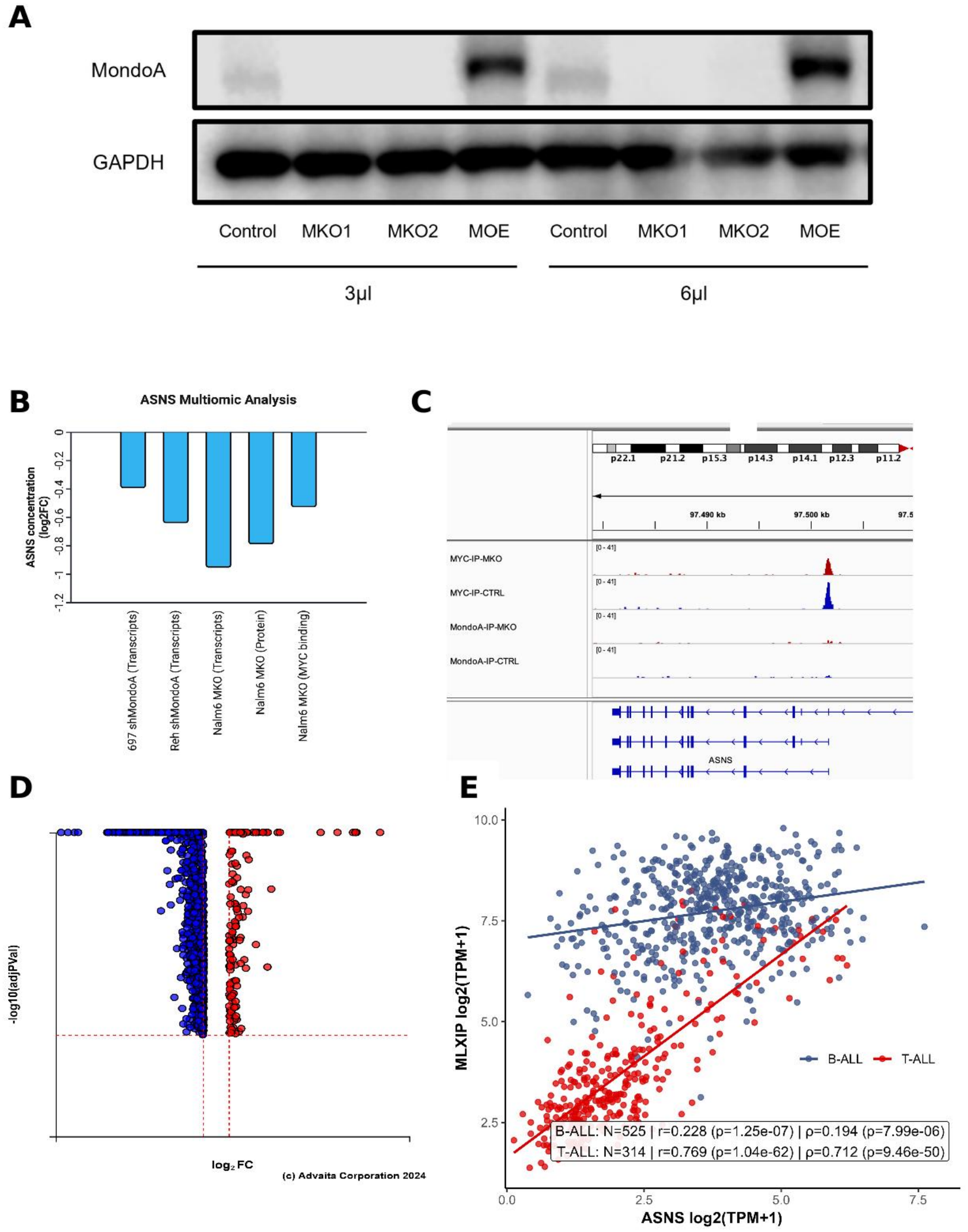
Validation of Nalm6 Models and Association of MondoA Status with ASNS Expression and MYC Binding. (A) MondoA Western blot in WT, MKO1, MKO2, and MOE Nalm6 clones, with GAPDH as loading control. The same lysates were loaded at 3 and 6 μL; the two loading volumes are not independent biological replicates. (B) ASNS mRNA changes in 697 shMondoA, REH shMondoA, and Nalm6 MKO (log2 fold changes: −0.39383, −0.64, and −0.9528, respectively), ASNS protein change in Nalm6 MKO2 (−0.789104), and MYC binding at the ASNS promoter. (C) MYC and MondoA ChIP-seq at the ASNS locus on chromosome 7: MYC signal is lower in MKO (red) than control cells (blue), with no clear MondoA promoter signal. (D) Genome-wide differential MYC promoter binding in MKO versus control cells (blue, decreased; red, increased). (E) ASNS and MondoA (MLXIP) transcript levels (log2[TPM+1]) with lineage-specific linear fits in inferred B-ALL (n = 525) and T-ALL (n = 314), reanalyzed from the UCSC Treehouse Tumor Compendium 25.01 PolyA.

### Cell viability experiments

Cell viability was measured using the RealTime-Glo™ MT Cell Viability Assay (Promega). Cells were seeded at 10,000 or 20,000 cells/well in 96-well plates containing Gln-replete or Gln-free RPMI 1640, with or without 10 U/mL L- asparaginase. Luminescence was monitored every 12 h for up to 84 h; cell-free medium wells served as background controls. GSH experiments used reduced glutathione supplementation in nutrient-replete, ASNase-containing, or Gln-free medium, as indicated in Figure 5. Comparator-drug experiments used 25 nM doxorubicin for 12 h or vincristine for 24 h. After background subtraction, time-course values were normalized to the corresponding luminescence at 0 h (100%). Endpoint treatment comparisons were normalized to the untreated control of the same genotype at the same time point, as specified in the figure legends. ASNase/control, combined-deprivation/ASNase, and GSH-associated viability ratios were calculated from these measurements. For Figure 5C–D and H, ratios were calculated for corresponding replicate wells before summarization.

### Flow Cytometry Apoptosis Assay (L-Asparaginase titration)

WT and MKO1 cells were seeded in triplicate in 24-well plates (1 × 10^6^ cells/well) and treated with ASNase at 0, 0.25, 0.5, 1, 5, or 10 U/mL. After 24, 48, or 72 h, cells were transferred to 96-well plates, centrifuged, washed with PBS, incubated with Annexin V/propidium iodide (PI) apoptosis assay reagents (Thermo Fisher) for 15 min, resuspended in Annexin binding buffer, and kept on ice. Flow cytometry was performed on a BD FACSCanto™ II and analyzed with FlowJo™ v10.8. Annexin V-positive/PI-negative cells were classified as early apoptotic and Annexin V-positive/PI-positive cells as late apoptotic. Figure 2A shows the 48 h endpoint.

**Figure 2:**
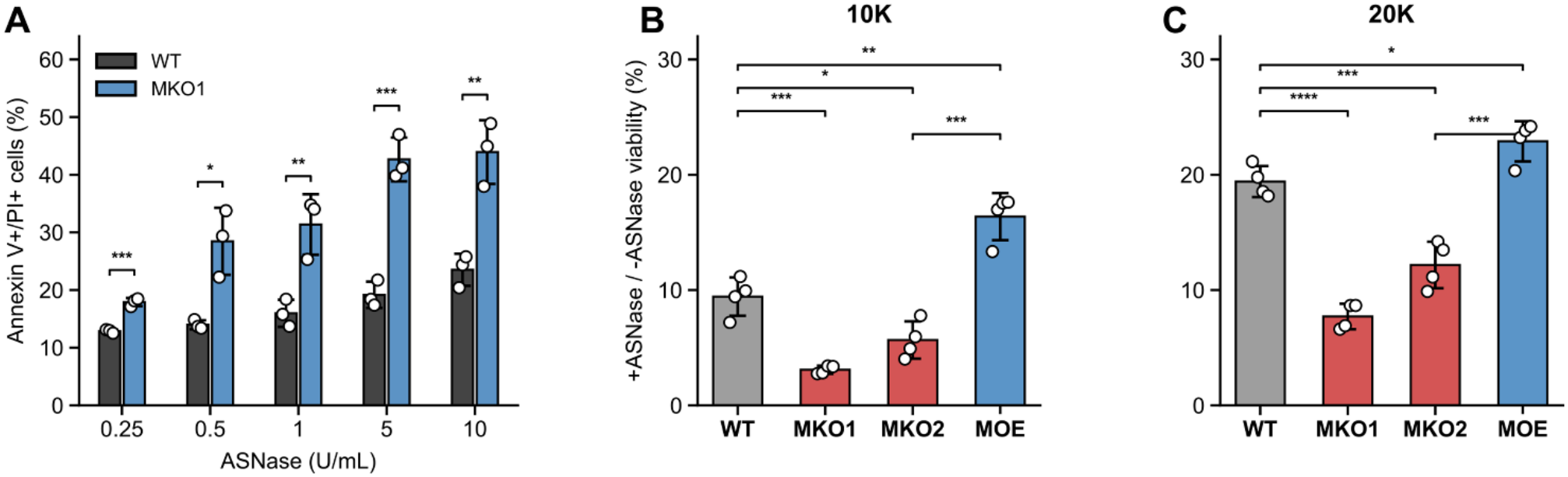
MondoA-Deficient B-ALL Cells Show Increased Sensitivity to L-Asparaginase. (A) Annexin V-positive/PI-positive late apoptotic WT and MKO1 Nalm6 cells after 48 h of treatment with 0.25–10 U/mL ASNase. (B–C) Viability after 72 h with 10 U/mL ASNase divided by viability in the corresponding untreated control, expressed as a percentage, at 10,000 (B) or 20,000 (C) cells/well. WT, MKO1, MKO2, and MOE clones were compared.

### Bioinformatic Analysis

Bulk RNA-seq gene-level expression (log2[TPM+1]) was obtained from the UCSC Treehouse Tumor Compendium 25.01 PolyA (GSE294351). ALL samples were retained. Because lineage annotation was unavailable, B-ALL and T-ALL were inferred using gene-signature scores: each marker gene was z-scored across ALL samples, and samples were classified according to the higher mean B-cell or T-cell marker score; samples with |B−T| < 0.25 were labeled ambiguous. B-cell markers were CD19, PAX5, EBF1, and IGLL1; T-cell markers were CD3D, CD3E, CD7, and LCK. Associations between ASNS and MLXIP (encoding MondoA) transcript levels were quantified separately within the inferred B-ALL and T-ALL groups using Pearson’s r and Spearman’s ρ with two-sided cor.test in R 4.4.2. Plots were generated with ggplot2; ambiguous samples were excluded from the displayed lineage-specific analyses.

### Multiomic analysis

Published Nalm6 expression-microarray (GSE168865) and MYC/MondoA ChIP-seq (GSE168864) datasets, together with transcriptomic comparisons of MondoA- depleted REH and 697 cells, were analyzed as previously described.(7) Mass- spectrometry-based proteomics was performed at BayBioMS@MRI. Cell pellets were lysed in urea extraction buffer, sonicated, clarified by centrifugation, and quantified by Bradford assay. After reduction, alkylation, and tryptic digestion, peptides were TMT10plex-labeled. High-pH reversed-phase fractionation generated 96 fractions, concatenated into 24 pools, acidified, and dried before LC–MS/MS. Nanoflow LC– MS/MS was performed on a Dionex 3000 coupled to an Orbitrap Eclipse using DDA with an MS3 method. Searches against the human reference proteome (UP000005640) and contaminants were performed in MaxQuant v2.3.0.0 with standard reporter-ion MS3 settings. Proteomics data were deposited to ProteomeXchange via PRIDE under PXD059106. Pathway and gene set enrichment analyses were conducted using iPathwayGuide.(9, 10)

### Western blotting

For Western blotting, 3 × 10^6^ cells were harvested and lysed in 50 μL RIPA buffer (20-188, Sigma) supplemented with protease inhibitors. Lysates were mixed with loading buffer and denatured at 70 °C for 10 min. For the clone-validation blot (Figure 1A), 3 or 6 μL of each lysate was loaded; for the treatment blots (Figure 3C), 8–15 μL was loaded. Proteins were resolved on 4%–12% SDS-PAGE gels (Thermo Fisher Scientific) and transferred onto 0.45 μm nitrocellulose membranes using the Trans-Blot Turbo system (Bio-Rad). Membranes were blocked with 5% BSA for 1 h at room temperature and incubated overnight at 4 °C with the following primary antibodies diluted 1:1,000: anti-MondoA polyclonal antibody (13614-1- AP, Proteintech), anti-PARP1 (9542S, Cell Signaling Technology), anti-GAPDH (3683S, Cell Signaling Technology), and anti-Actin (66009-1-Ig, Proteintech). The clone-validation blot was probed for MondoA and GAPDH; the treatment blots were probed for PARP1, MondoA, GAPDH, and Actin. GAPDH and Actin served as loading controls. After washing, membranes were incubated with the appropriate secondary antibodies, and signals were detected using an enhanced chemiluminescence substrate. PARP1 cleavage was summarized as the densitometric ratio of cleaved to full-length PARP1 (Figure 3D).

**Figure 3:**
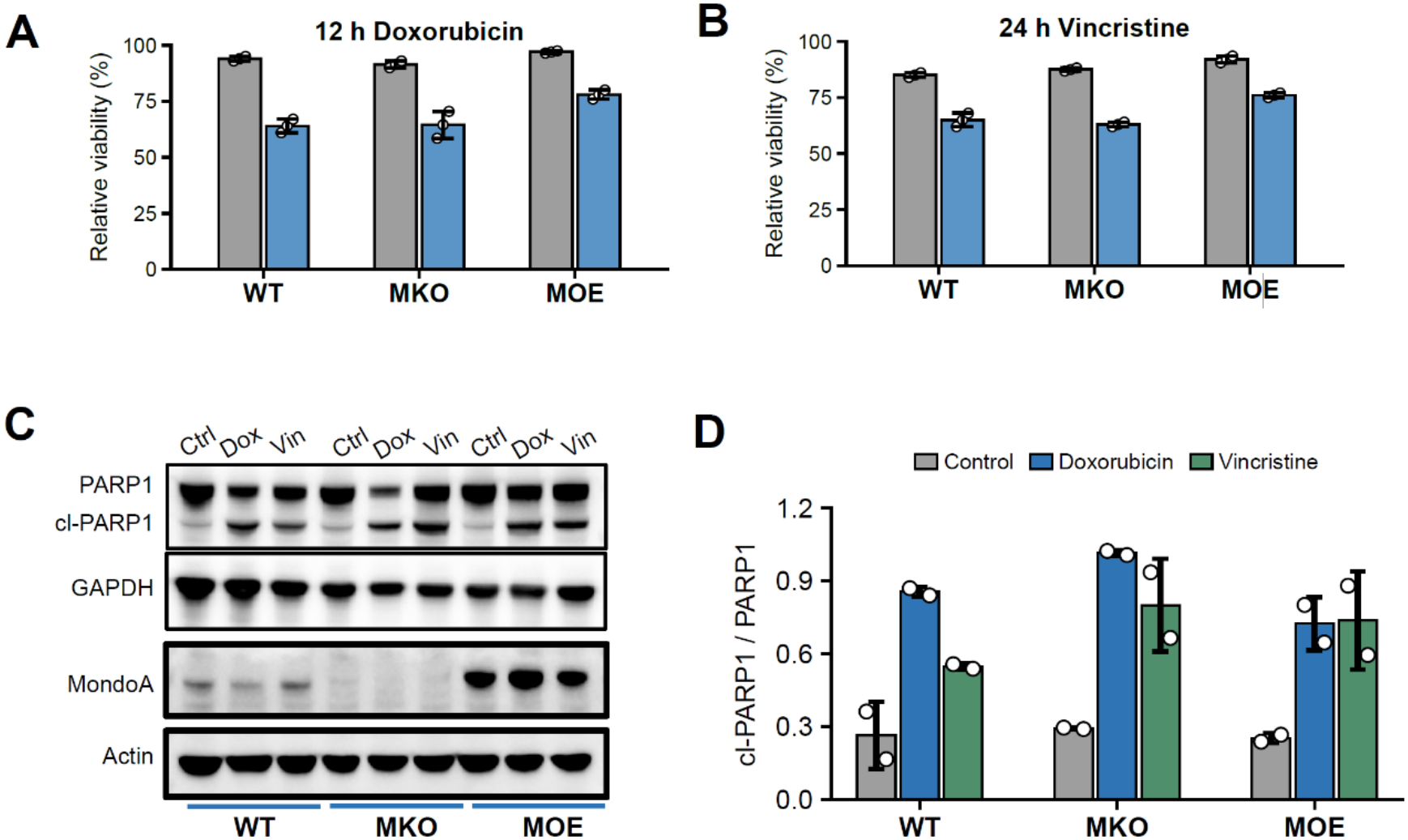
MondoA Deficiency Does Not Increase Sensitivity to Doxorubicin or Vincristine. (A–B) Relative viability of WT, MKO1, and MOE Nalm6 cells after 12 h with 0 or 25 nM doxorubicin (A) or after 24 h with or without vincristine (B). (C) Western blots of full-length PARP1, cleaved PARP1 (cl- PARP1), GAPDH, MondoA, and Actin in untreated (Ctrl) or drug-treated cells. GAPDH and Actin served as loading controls. (D) Densitometric ratio of cleaved to full-length PARP1. MKO denotes MKO1 throughout this figure.

### Statistical analysis

P values were calculated using unpaired, two-sided Student’s t-tests assuming equal variances. Figure 5 P values were unadjusted; P < 0.05 was considered significant.

### Funding

This work was supported by Cura Placida – Children’s Cancer Research Foundation.

## Results

### MondoA Status Is Associated with ASNS Expression and MYC Binding at the ASNS Promoter

We first verified MondoA expression in the Nalm6 models. Western blotting showed markedly reduced MondoA protein in MKO1 and MKO2 and increased expression in MOE relative to WT (Figure 1A). We then integrated transcriptomic, proteomic, and ChIP-seq data to examine the relationship between MondoA and ASNS. Published expression-microarray comparisons included Nalm6 MKO versus control cells and REH and 697 cells expressing shMondoA versus control shRNA.(7) ASNS mRNA was lower in each MondoA-deficient condition, and proteomics showed lower ASNS protein abundance in MKO2 cells (Figure 1B). MYC occupancy at the ASNS promoter was reduced in MKO cells, whereas no clear MondoA occupancy was detected at this locus (Figure 1C). Together with the previously reported genome- wide redistribution of MYC binding after MondoA loss (Figure 1D), these findings are consistent with an indirect link between MondoA status and ASNS transcription. They do not establish that altered MYC occupancy causes the change in ASNS expression. We additionally reanalyzed ALL RNA-seq data from the UCSC Treehouse Tumor Compendium 25.01 PolyA (GSE294351). MondoA and ASNS transcript levels were positively correlated in the inferred B-ALL group (n = 525; r = 0.228; P = 1.25 × 10−7) and more strongly correlated in the inferred T-ALL group (n = 314; r = 0.769; P = 1.04 × 10−62; Figure 1E). These patient-data associations support a relationship between MondoA and ASNS expression beyond the Nalm6 model, but do not establish treatment resistance in either lineage.

### MondoA-Deficient B-ALL Cells Show Increased Sensitivity to L-Asparaginase

Using the WT, MKO1, MKO2, and MOE Nalm6 clones defined above, we next analyzed the effect of MondoA status on ASNase sensitivity. The apoptosis titration compared WT with MKO1 only; MKO2 was evaluated in the complementary viability experiments. After 48 h of treatment with 0.25–10 U/mL ASNase, Annexin V-positive/PI-positive late apoptotic cells were significantly more frequent in MKO1 than in WT at all tested concentrations (Figure 2A). To complement the apoptosis analysis, we examined ASNase effects on viability in all four clones. ASNase reduced viability in WT, MKO1, MKO2, and MOE cells (Supplementary Figure 1A–B). We then divided viability under ASNase treatment by viability in the corresponding untreated control. This ratio was higher in WT and MOE than in the knockout clones at both seeding densities, indicating greater relative preservation of viability in MondoA-expressing cells (Figure 2B–C). Together, the apoptosis and viability data show increased ASNase sensitivity after MondoA loss.

### MondoA Deficiency Does Not Increase Sensitivity to Doxorubicin or Vincristine

To determine whether the increased ASNase sensitivity reflected a general vulnerability to cytotoxic agents, we exposed WT, MKO1, and MOE Nalm6 clones to doxorubicin (Dox) or vincristine (Vin). These comparator experiments used MKO1 only. Dox and Vin reduced viability in all tested clones, with no evidence of increased sensitivity in MKO1 relative to WT under these conditions (Figure 3A–B). We also assessed apoptosis by measuring cleaved PARP1 and the cleaved-to-full-length PARP1 ratio. These measurements showed no consistent increase in drug-induced PARP1 cleavage in MKO1 relative to WT or MOE (Figure 3C–D). Thus, the enhanced ASNase response associated with MondoA loss was not accompanied by a comparable increase in sensitivity to Dox or Vin in the tested MKO1 clone.

### Glutamine Withdrawal Reduces the Viability Advantage of MondoA-Proficient Cells during Asparaginase Treatment

We examined whether combining ASNase treatment with Gln withdrawal increased cytotoxicity and whether this additional effect differed according to MondoA status. Nalm6 clones were cultured for 72 h in Gln-replete control medium, Gln-free medium, ASNase- containing medium, or Gln-free medium containing ASNase. Gln withdrawal reduced viability relative to control medium, and combined treatment further reduced viability compared with Gln withdrawal alone (Figure 4A). At 10,000 cells/well, the ratio of viability under combined treatment to viability under ASNase alone was significantly higher in MKO1 (p< 0.01) and MKO2 (p<0.05) compared toWT (), indicating a smaller additional effect of Gln withdrawal in the MKO condition; (Figure 4B). The magnitude of the additional viability loss was smaller in the knockout clones, particularly MKO1, than in WT and MOE at both seeding densities (Figure 4C). Thus, Gln withdrawal diminished the relative preservation of viability in MondoA- proficient clones during ASNase treatment under the condition tested. This pattern is compatible with greater dependence on Gln-supported Asn synthesis in MondoA-proficient cells, although ASNS activity was not directly measured.

**Figure 4:**
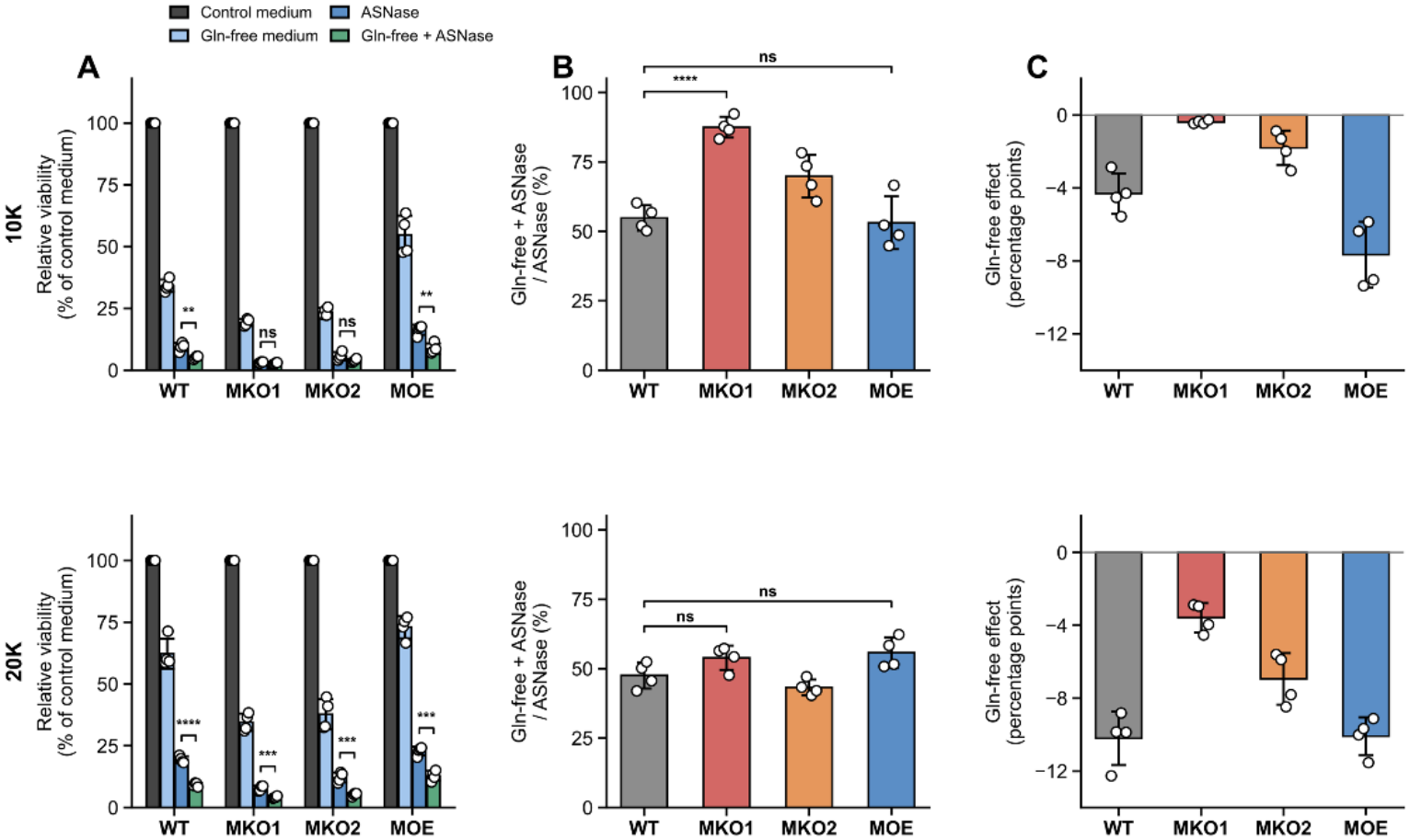
Glutamine Withdrawal Reduces the Viability Advantage of MondoA-Proficient Cells during Asparaginase Treatment. (A) Viability after 72 h in control medium, Gln-free medium, 10 U/mL ASNase, or Gln-free medium with 10 U/mL ASNase, normalized to the corresponding untreated control. (B) Viability in Gln-free medium with ASNase divided by viability in Gln-replete medium with ASNase, expressed as a percentage. (C) Change in viability upon Gln withdrawal during ASNase treatment, calculated as viability in Gln-free medium with ASNase minus viability in Gln-replete medium with ASNase, in percentage points; negative values indicate an additional loss of viability. Upper and lower rows show 10,000 and 20,000 cells/well, respectively.

### Reduced Glutathione Increases Viability in MondoA-Knockout Clones Particularly Under Amide Amino Acid Deprivation

Our previous work linked MondoA loss to increased mitochondrial respiration and impaired control of ROS during Gln starvation.(7) We therefore tested reduced glutathione (GSH), whose synthesis uses glutamate derived from Gln, to examine whether GSH supplementation could preserve viability during amino acid stress. At 72 h in nutrient-replete RPMI 1640 (10,000 cells/well), GSH decreased viability in MKO1 (P = 0.00013), whereas WT and MKO2 showed no significant change (Figure 5A). During ASNase treatment, mean viability increased with GSH in WT, MKO1, and MKO2, but this increase was significant only in MKO2 (Figure 5B). GSH-associated fold increase during ASNase treatment was significantly greater in MKO2 than in WT; MKO1 showed the same trend without a significant difference from WT (Figure 5C–D). Thus, we used MKO2 in the following experiments. In GSH-supplemented nutrient-replete medium, MKO2 viability exceeded WT from 36 h onward (Figure 5E). At 72 h, the MKO2-to-WT viability ratio was greater under ASNase treatment or Gln withdrawal than in nutrient-replete medium (Figure 5F–H). These findings extend our previous observations of impaired metabolic-stress adaptation after MondoA loss.(7) Together, they show GSH-associated preservation of viability during amino acid stress, most prominently in MKO2, without establishing the mechanism of this protection.

**Figure 5:**
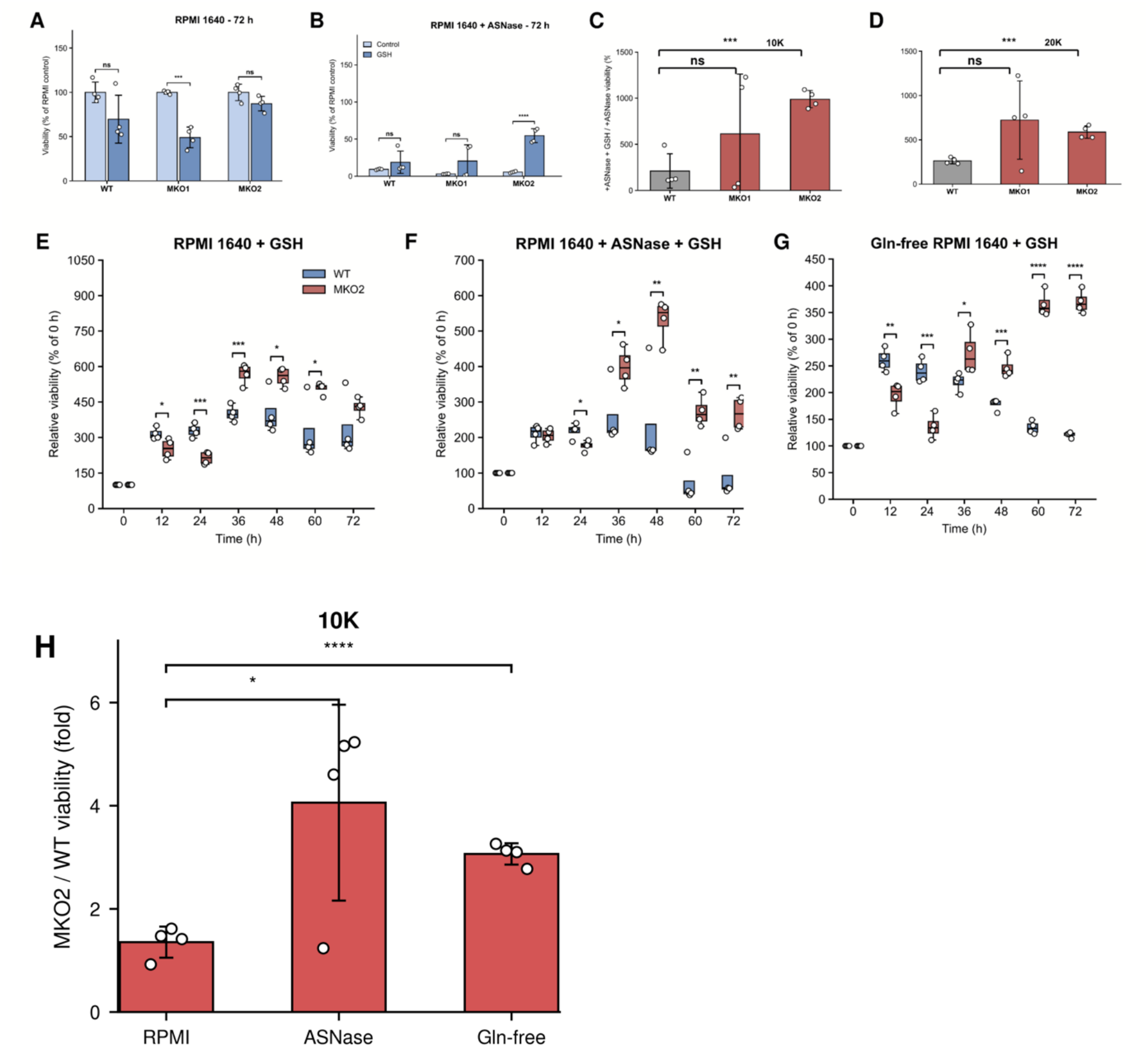
Reduced Glutathione Increases Viability in MondoA-Knockout Clones Particularly Under Amide Amino Acid Deprivation. (A–B) Relative viability of WT, MKO1 and MKO2 Nalm6 cells after 72 h in RPMI 1640 without or with GSH (A) or in RPMI 1640 with 10 U/mL ASNase, without or with GSH (B), normalized to the untreated RPMI 1640 control of the same genotype at 72 h. (C–D) GSH-associated fold change in viability during ASNase treatment, shown as viability with ASNase and GSH relative to viability with ASNase alone, at 10,000 (C) or 20,000 (D) cells per well, expressed as a percentage. (E–G) Relative viability of WT and MKO2 Nalm6 cells over 72 h in RPMI 1640 with GSH (E), RPMI 1640 with 10 U/mL ASNase and GSH (F), or Gln-free RPMI 1640 with GSH (G), normalized to the corresponding value at 0 h. (H) Relative viability of MKO2 normalized to WT at 72 h across the three GSH-containing media shown in E–G, at 10,000 cells per well; the ASNase-containing and Gln- free conditions were each compared with RPMI 1640. Cells were cultured at 10,000 cells per well unless otherwise indicated. Data represent four replicate wells per condition. For C–D and H, ratios were calculated for corresponding replicate wells before summarization. Unadjusted P values were calculated using unpaired, two-sided Student’s t-tests assuming equal variances. Statistical significance is indicated as follows: ns, P ≥ 0.05; *P < 0.05; **P < 0.01; ***P < 0.001; ****P < 0.0001.

## Discussion

ASNase is a cornerstone of ALL therapy, and resistance to this agent contributes to treatment failure and relapse. Here, MondoA-deficient Nalm6 B-ALL cells were more sensitive to ASNase than MondoA-proficient counterparts.

Mechanistically, MondoA deficiency was associated with lower ASNS mRNA in Nalm6, REH, and 697 cells and lower ASNS protein abundance in Nalm6 MKO2. Because ASNS is associated with ASNase resistance,(6) this provides a plausible link between MondoA status and treatment response. MYC occupancy at the ASNS promoter was also reduced in MondoA-knockout cells. No clear MondoA occupancy was detected at this locus in the analyzed ChIP-seq data, favoring an indirect model involving MYC or broader transcriptional changes. However, the absence of a detectable peak does not exclude direct regulation, and the causal contribution of altered MYC binding remains unresolved. This interpretation is consistent with our previous work showing that MondoA enables B-ALL cells to adapt to Gln starvation by restricting oxidative phosphorylation and limiting ROS production.

Combined ASNase treatment and Gln withdrawal resulted in the lowest viability compared with untreated control medium, ASNase treatment alone, and Gln withdrawal alone. Gln withdrawal caused a smaller additional reduction during ASNase exposure in the knockout clones, particularly MKO1, consistent with greater dependence on Gln-supported ASNS activity in MondoA-proficient cells. However, the low viability already reached with ASNase alone in MKO cells may also limit the measurable additional effect of Gln withdrawal. The GSH-associated viability gain relative to WT was most pronounced in MKO2. This difference cannot be attributed solely to monoallelic versus biallelic disruption on the basis of two clones; other clone-specific properties may contribute.

These data link MondoA status to ASNS abundance and adaptation to ASNase- induced amino acid stress. The Amino Acid Response (AAR) provides a relevant framework: Asn and Gln depletion can activate GCN2–eIF2α–ATF4 signaling, reduce global protein synthesis, and induce adaptive genes such as ASNS.(11, 12) We did not test activation of this signaling axis, so our findings do not establish that MondoA is part of the canonical GCN2–ATF4 pathway.

We titrated ASNase from 0.25 to 10 U/mL and used 10 U/mL for the viability experiments. This exceeds the commonly used clinical target trough activity of 100 U/L (0.1 U/mL).(13) The higher concentration was chosen to sustain amino acid depletion over the observation period. Because extracellular Asn and Gln concentrations were not measured, the extent and persistence of depletion were not directly verified. Responses at clinically relevant ASNase activities require further study.

GSH was selected because Gln supplies glutamate for its synthesis and our prior work implicated oxidative stress in MondoA-deficient cells. Other antioxidants were not compared, and ROS/mtROS, intracellular GSH, and amino acid flux were not measured during ASNase exposure. Thus, the viability response does not establish a ROS-dependent mechanism or preferential GSH utilization by MKO2. Mitochondrial GSH homeostasis can also support oxidative phosphorylation,(14) providing an additional explanation for GSH-associated effects on viability beyond antioxidant buffering.

ASNase sensitivity was evaluated in Nalm6-derived clones and was not tested in vivo. Although the transcriptomic association with ASNS extended to REH and 697, treatment responses require validation in independent leukemia models, including MondoA-depleted xenografts. Defining the MondoA–ASNS relationship will also require functional ASNS rescue and targeted analysis of MYC-dependent regulation at the ASNS locus. The available ChIP-seq and expression data support this model but do not establish a direct regulatory mechanism.

Together, these in vitro findings associate MondoA deficiency with lower ASNS abundance and greater ASNase sensitivity in Nalm6 cells, and identify GSH- associated protection during amino acid stress. Further mechanistic and in vivo validation is required before MondoA can be established as a biomarker or therapeutic target for ASNase response.

## Declaration of generative AI and AI-assisted technologies in the manuscript preparation process

During the preparation of this work, the authors used OpenAI ChatGPT (Codex) to improve language, clarity and consistency in the revised manuscript text. After using this tool, the authors reviewed and edited the content as needed and take full responsibility for the content of the published article.

---

**Supplementary Figure 1: MondoA-Deficient B-ALL Cells Show Increased Sensitivity to L-Asparaginase**. Viability of WT, MKO1, MKO2, and MOE Nalm6 cells after 72 h with or without 10 U/mL ASNase, normalized to the corresponding untreated control (100%), at 10,000 (A) or 20,000 (B) cells/well.

