## Supplementary figures and images for "Targeting MondoA Sensitizes B-ALL Cells to Asparaginase and Reveals Glutathione-Associated Protection"

### Supplementary Figure 1

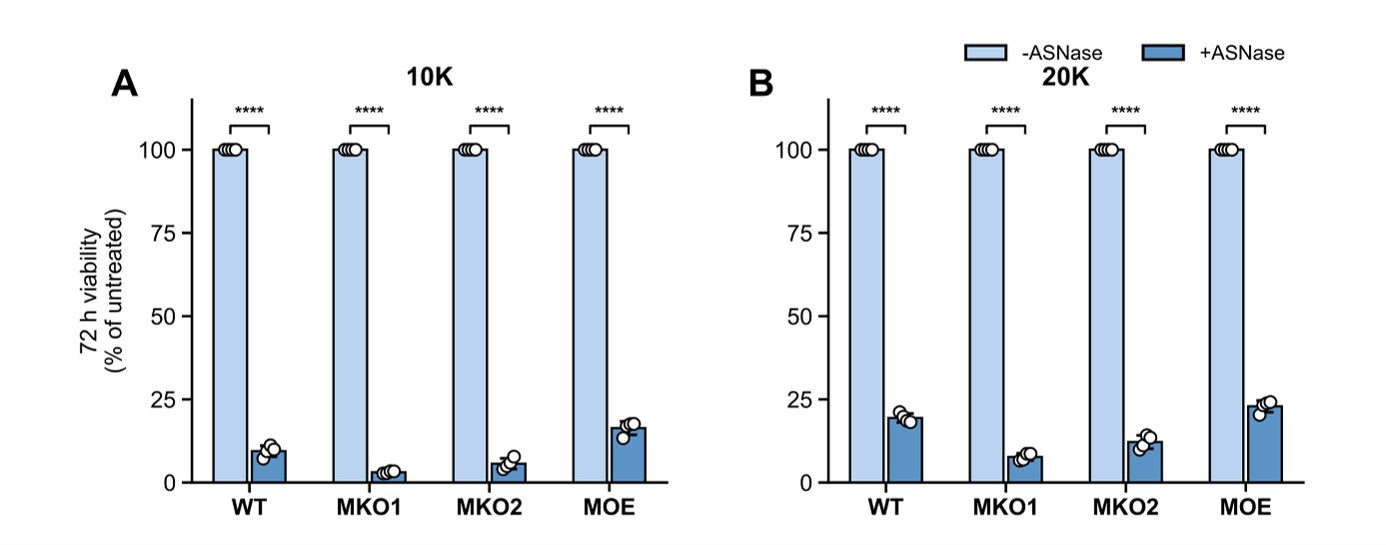
